# Whole genome sequence analysis of 91 *Salmonella* Enteritidis isolates from mice caught on poultry farms in the mid 1990s

**DOI:** 10.1101/301432

**Authors:** Jean Guard, Guojie Cao, Yan Luo, Joseph D. Baugher, Sherrill Davison, Kuan Yao, Maria Hoffmann, Guodong Zhang, Rebecca Bell, Jie Zheng, Eric Brown, Marc Allard

## Abstract

*Salmonella enterica* serovar Enteritidis (SE), the most commonly reported serovar of human salmonellosis, has been frequently associated with poultry farms, eggs and egg products. Mice are known vectors of SE contamination in these facilities. The objective of this study was to use whole-genome sequencing (WGS) to analyze SE from mice obtained at poultry farms in Pennsylvania. Documenting pathogen diversity can identify reliable biomarkers for rapid detection and speed up outbreak investigations. We sequenced 91 SE isolates from 83 mice (62 spleen isolates, 29 intestinal isolates) caught at 15 poultry farms between 1995-1998 using an Illumina NextSeq 500. We identified 742 single nucleotide polymorphisms (SNPs) capable of distinguishing each isolate from one another. Isolates were divided into two major clades: there were more SNPs differences within Clade B than counterparts in Clade A. All isolates containing antimicrobial resistance genes belong to Subgroup B2. Clade-defining SNPs provided biomarkers distinguishing isolates from 12 individual subgroups, which were separated by farm location or year of collection. Nonsynonymous changes from the clade-defining SNPs proffered a better understanding of possible genetic variations among these isolates. For a broader view of SE diversity, we included data from NCBI Pathogen Detection Isolates Browser, in which subgroups in Clade B formed new SNP Clusters.

**Importance:** WGS and SNPs analyses are excellent and powerful tools for investigating SE phylogenies. Identifying the evolutionary relationships among SE isolates from mouse, poultry, environmental, and clinical isolates, along with patterns of genetic diversity, advances understanding of SE and the role mice may play in SE contamination and spread among poultry population. Our data was able to identify SE isolates from different farms or years of collection. Moreover, the annotations of clade-defining SNPs provided information about possible protein functions among these SE isolates from each subgroup. Clade-defining or farm-unique biomarkers were useful for rapid detection and outbreak investigations.

## Introduction

*Salmonella enterica* serovar Enteritidis (SE) is a long-standing public health concern in the US (1); salmonellosis can result in hospitalization or death of infants, the elderly, and those with compromised immune systems (2, 3). This pathogen has been strongly associated with poultry farms, eggs, and egg products (4, 5). In 2010, SE linked to shell eggs resulted in an outbreak requiring the recall of a half billion eggs (https://www.cdc.gov/salmonella/2010/shell-eggs-12-2-10.html) (6).

One of the challenges in resolving foodborne outbreaks associated with SE is the extreme genomic homogeneity within a specific geographic location or ecology system and its broad host range (6, 7). Mice are important biological vehicles contributing to SE dissemination and amplification in chicken houses, especially among laying hens (8, 9). In fact, SE has been strongly correlated with rodent activity; chickens in caged housing where mice are present are more likely to carry SE (10). Understanding the evolutionary relationships among SE isolates from mice, poultry, environmental surfaces, and clinical cases is important both for outbreak investigations and for identifying strains with genetic markers for virulence or capacity for rapid host adaptation, such as mutations in the mismatch repair gene *mutS* that can contribute to rapid evolution in immunocompromised hosts (11).

Whole genome sequencing (WGS) methods have identified variations across otherwise indistinguishable isolates from eggs and egg products (6, 12), SE associated with reptile feeder mice (13), S. Montevideo from red and black pepper (14). Genome-wide single nucleotide polymorphisms (SNPs) detected by WGS are considered as the most valuable genetic markers for investigating the evolutionary relationships among SE homogeneous isolates (1, 7, 15). Application of WGS have also been useful in other microorganisms, including *E. coli* (16), *Vibrio cholera* (17), and *Staphylococcus aureus* (18).

Importantly, WGS can be also applied to historic isolates, some of which have been stored for decades. Data from those historic isolates should allow us to understand the origin and persistence of important traits. In this current project, we sequenced 91 SE isolated from 82 mice at poultry farms during the 1990s, which lets us to compare both site and host-adaptions with those of isolates from more recent sampling. Documenting these genomes and fitting them into large-scale phylogeny projects such as GenomeTrakr (https://www.fda.gov/Food/FoodScienceResearch/WholeGenomeSequencingProgramWGS/ucm363134.htm) and NCBI Pathogen Detection Isolates Browser (https://www.ncbi.nlm.nih.gov/pathogens/) will refine our understanding of SE contamination and spread in poultry facilities (19). Further, identifying and characterizing biomarkers can facilitate the development of rapid and reliable tests that could guide appropriate interventions during future outbreaks.

## Materials and Methods

### Bacterial isolates

Ninety-one SE isolates from mouse spleens (n=62) and intestines (n=29), collected from 15 poultry farms in Pennsylvania during 1995-1998, are listed in Table 1. Among these isolates, eight pairs were isolated from the spleen and intestine of the same mouse; these were designated as m1 through m8. These isolates are archived under Bioproject Number PRJNA186035 (https://www.ncbi.nlm.nih.gov/bioproject/186035).

**Table 1.**
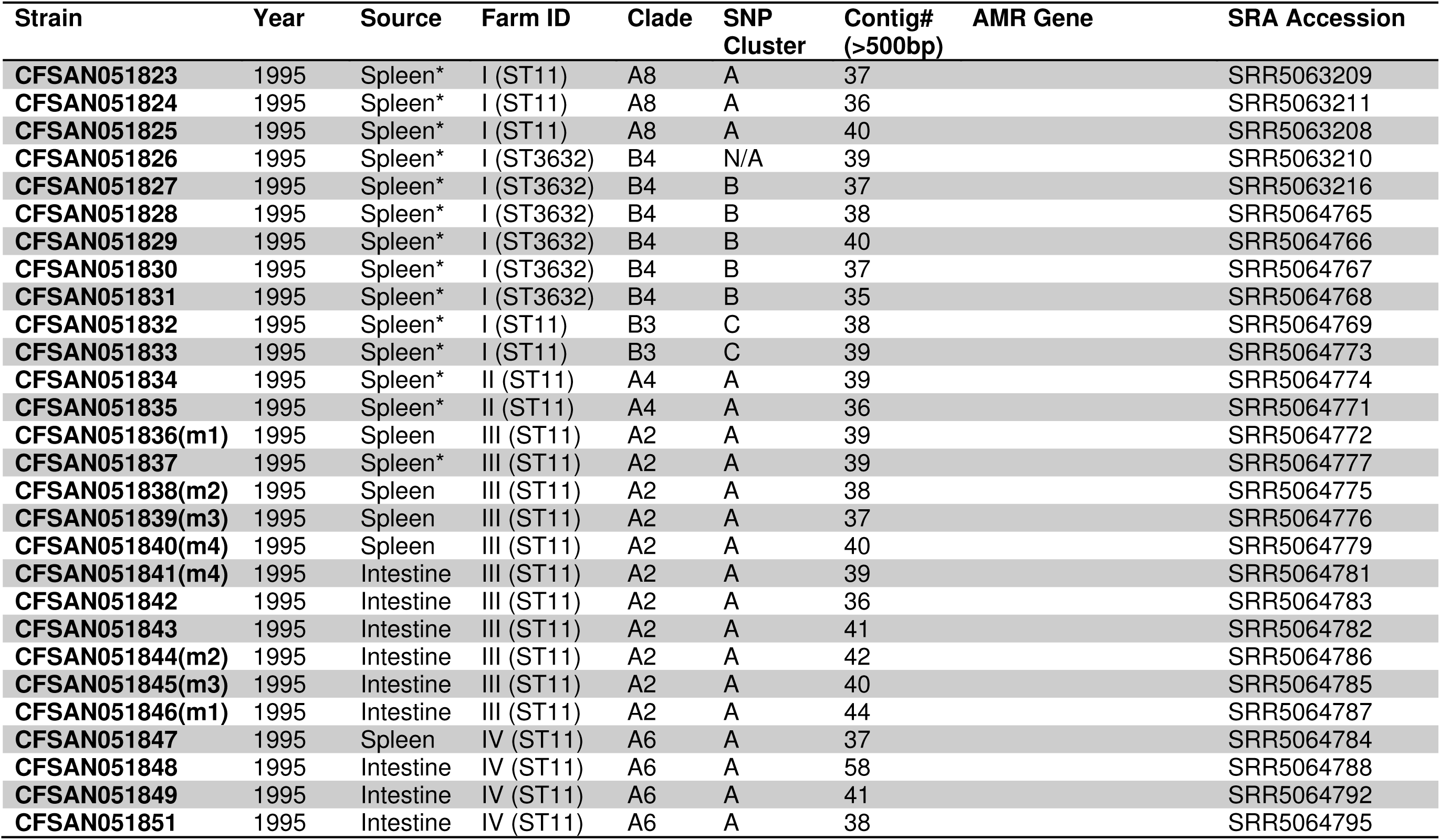
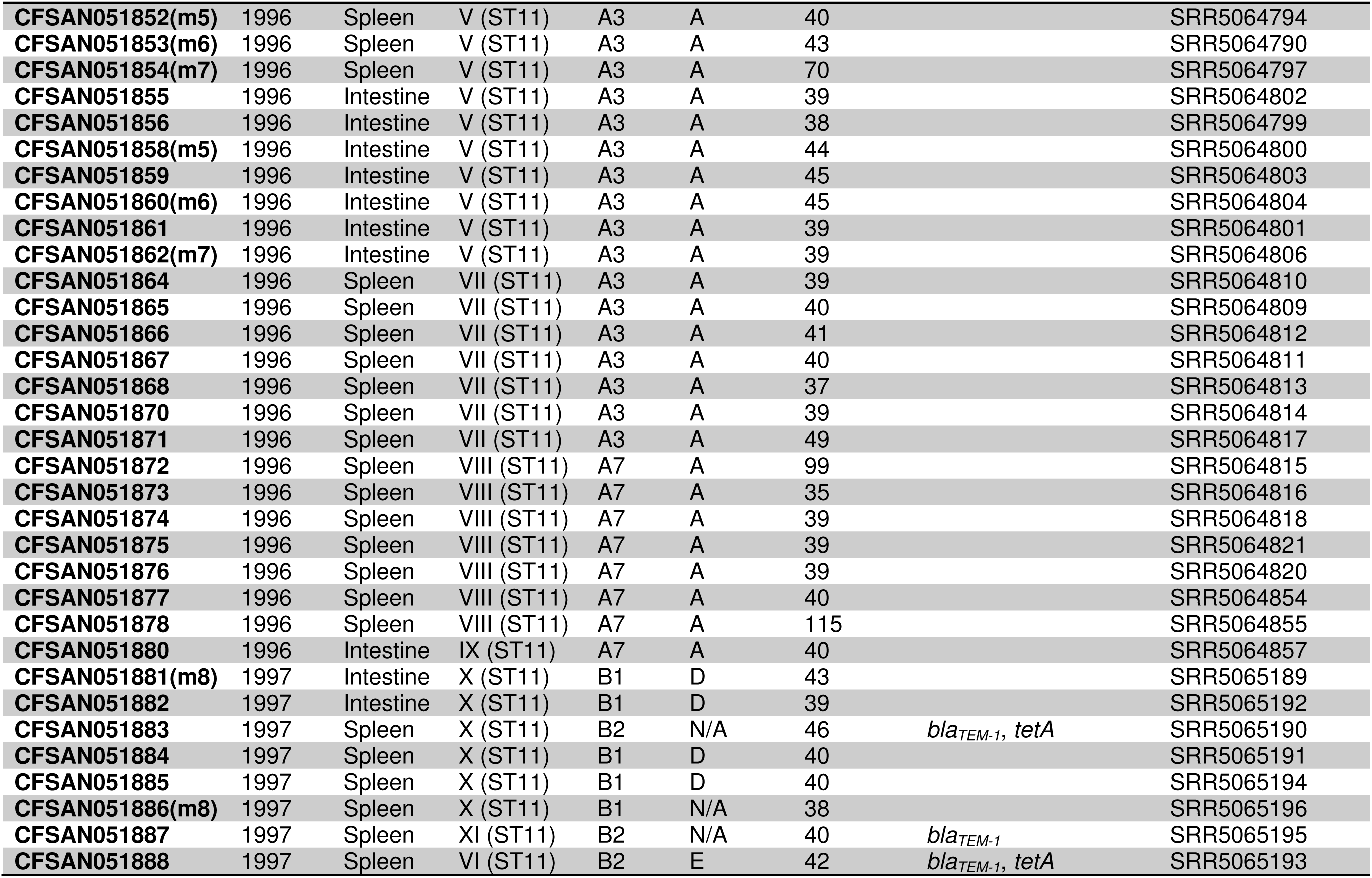
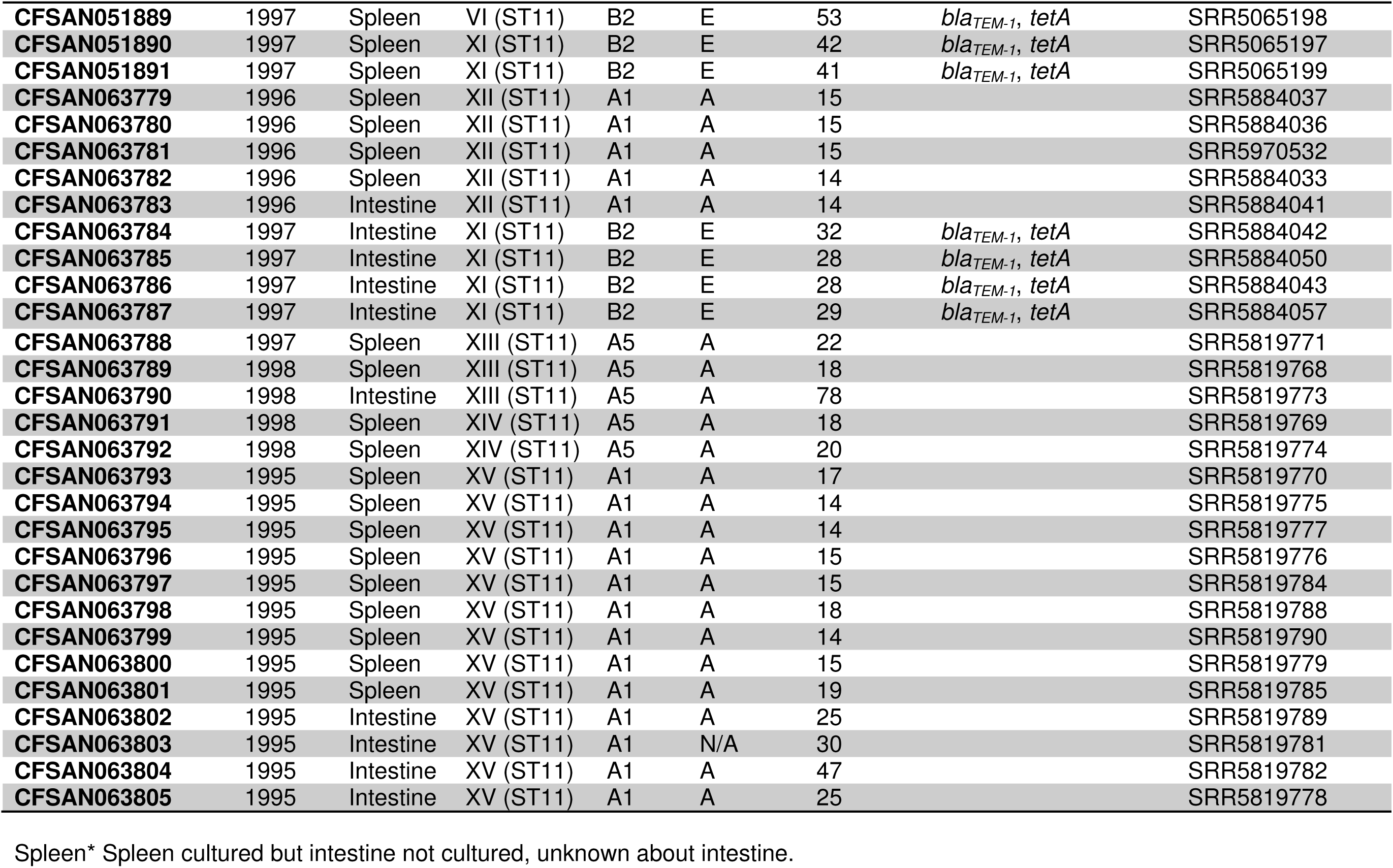
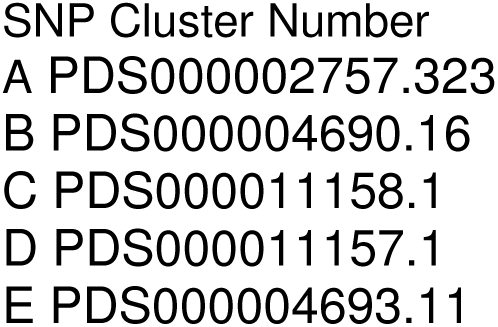
The metadata and general genomic information of 91 sequenced S. Enteritidis in current study.

**Table 2.** The number of SNP differences (standard deviation) between 12 subgroups.

|  | A1 | A2 | A3 | A4 | A5 | A6 | A7 | A8 | B1 | B2 | B3 |
| --- | --- | --- | --- | --- | --- | --- | --- | --- | --- | --- | --- |
| **A2** | 44(5) |  |  |  |  |  |  |  |  |  |  |
| **A3** | 50(6) | 45(6) |  |  |  |  |  |  |  |  |  |
| **A4** | 40(5) | 36(5) | 39(6) |  |  |  |  |  |  |  |  |
| **A5** | 44(5) | 40(5) | 43(6) | 32(5) |  |  |  |  |  |  |  |
| **A6** | 54(6) | 50(6) | 55(7) | 46(6) | 50(6) |  |  |  |  |  |  |
| **A7** | 51(6) | 47(6) | 52(6) | 43(6) | 47(6) | 57(7) |  |  |  |  |  |
| **A8** | 50(6) | 45(6) | 51(6) | 41(6) | 45(6) | 55(7) | 24(5) |  |  |  |  |
| **B1** | 92(8) | 90(8) | 95(8) | 86(8) | 90(8) | 99(8) | 97(8) | 95(8) |  |  |  |
| **B2** | 84(8) | 82(8) | 87(8) | 78(8) | 82(8) | 91(8) | 89(8) | 87(8) | 33(5) |  |  |
| **B3** | 137(9) | 135(9) | 140(10) | 131(9) | 135(9) | 144(10) | 142(10) | 140(10) | 86(8) | 78(8) |  |
| **B4** | 133(9) | 131(9) | 137(9) | 127(9) | 132(9) | 141(10) | 138(10) | 137(9) | 83(8) | 75(7) | 46(6) |

### Whole genome sequencing and assembly

Genomic DNA was extracted after incubation of culture for 16 hours at 37 °C in Trypticase Soy Broth (TSB) using the DNeasy Blood and Tissue Kit (Qiagen Inc, Valencia, CA). Concentrations of DNA were measured using a Qubit 3.0 fluorometer (Life Technologies, MD). Libraries were prepared according to Nextera XT protocols and sequenced on the Illumina NextSeq 500 (Illumina, San Diego, CA) using NextSeq 500/550 High Output Kit v2 (300 cycles). Raw reads were assembled *de novo* using SPAdes software v3.8.2 with default settings (20). We obtained chromosome draft genomes between 4.69M bps and 4.80M bps. These genomes were annotated using the NCBI Prokaryotic Genome Annotation Pipeline (PGAP) (21).

We selected SE CFSAN051873 (spleen, 1996, Farm VIII) to serve as the reference genome, using the PacBio platform we obtained a fully closed genome for CFSAN051873 as follows (22). Genomic DNA was sheared into approximately 20-kb fragments using g-TUBE (Covaris, Inc., Woburn, MA). The library was prepared based on the 20-kb PacBio sample preparation protocol and sequenced using P6/C4 chemistry on four single-molecule real-time (SMRT) cells with a 240-min collection time. The continuous long-read data were *de novo* assembled using the PacBio hierarchical genome assembly process (HGAP version 3.0) with default parameters (23). The assembled sequence was annotated using PGAP (21).

### Genomic and phylogenetic analysis

The Fastq data from NextSeq runs were put into the Center for Food Safety and Applied Nutrition (CFSAN) SNP pipeline v0.8 to create a SNP matrix (24) with SE CFSAN051873 (CP_022003.1) as the reference genome. GARLI (Genetic Algorithm for Rapid Likelihood Inference: https://code.google.com/archive/p/garli/) v2.01 (25) was used to construct maximum-likelihood (ML) phylogenetic trees (ratematrix = 6rate; ratehetmodel = gamma). Multiple runs were performed (n=100) to ensure that results were consistent. To estimate support for each node, phylogenies were created for 1,000 bootstrap replicates of the data set from GARLI. Python program SumTrees was used to generate one consensus tree with bootstrap values at a 70% threshold (https://pvthonhosted.org/DendroPv/programs/sumtrees.html) and FigTree v 1.4.3 was used to export the figures (http://tree.bio.ed.ac.uk/software/figtree/). NCBI Pathogen Detection Isolates Browser (https://www.ncbi.nlm.nih.gov/pathogens) was used to show phylogenetic relationship among SE isolates from broader ranges of geographical locations and sources. Custom script was used to identify clade-defining SNPs and Tool for Rapid Annotation of Microbial SNPs (TRAMS) tool to perform annotations on clade-defining SNPs (26). The pairwise distance matrix, shown as number of SNP differences among isolates, was calculated using MEGA7 with 1,000 bootstrap iterations (27).

## Results

### Phylogenetic analysis

#### Overview

We identified 742 SNPs and generated the maximum-likelihood phylogenetic tree arising from these SNPs, as depicted in Fig 1. Tree tips were marked using isolate name, source, year, farm, and NCBI Pathogen Detection Isolates Browser SNP Cluster. For example, CFSAN051866 was labeled as CFSAN051866_spleen_1996_FarmVII_SCA, which provides the following details: this bacterium was isolated from a mouse spleen in 1996, that mouse came from Farm VII, and the isolate fits within SNP Cluster A (28), which was designated according to the NCBI Pathogen Detection Isolates Browser (Table 1). Subgroup names and the number of clade-defining SNPs were labeled on the internal branches. For example, Subgroup B1 had the most clade-defining SNPs (179 SNPs), while Subgroup A5 had only 6 clade-defining SNPs.

**Figure 1.**
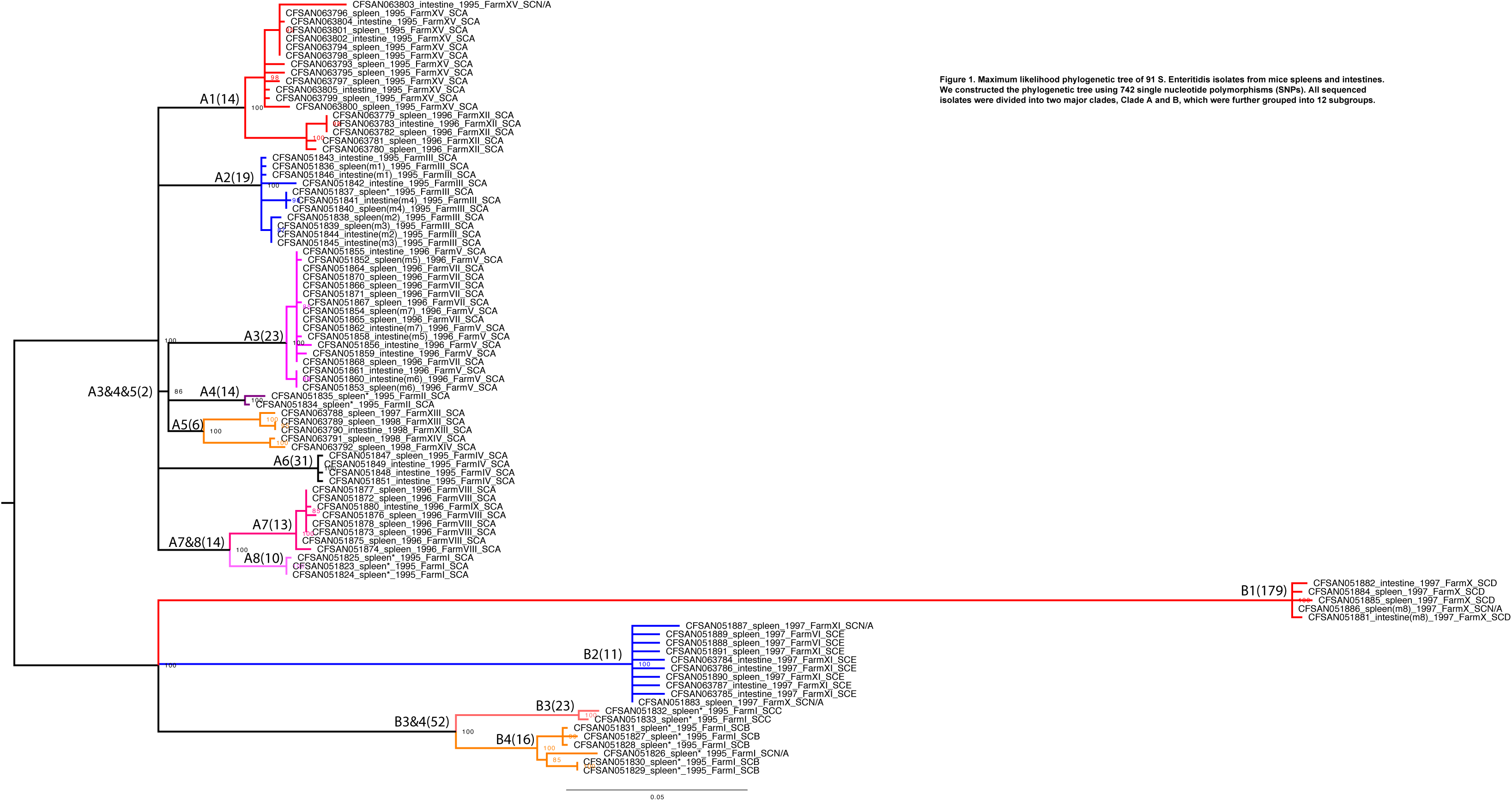
Maximum likelihood phylogenetic tree of 91 S. Enteritidis isolates from mice spleens and intestines. We constructed the phylogenetic tree using 742 single nucleotide polymorphisms (SNPs). All sequenced isolates were divided into two major clades, Clade A and B, which were further grouped into 12 subgroups.

#### Phylogenetic Tree Construction

We recognized two major clades: Clades A and B, which further subdivided into 12 subgroups: A1 to A8 and B1 to B4. It was notable that all isolates carrying antimicrobial resistance genes belonged exclusively to Subgroup B2. Moreover, isolates in each subgroup had varied ranges of SNP differences. The maximum SNP differences within Subgroups A1 and A5 were 33 (CFSAN063779 and CFSAN063803) and 27 (CFSAN063788 and CFSAN063792) SNPs, respectively, while the maximum number in Subgroup A3 was only 6 SNPs (CFSAN051856 and CFSAN051861). Subgroups A1 and A2 were the two largest subgroups, containing 18 and 17 isolates, respectively. Subgroup B3 only contained two isolates, as the smallest group in the tree.

#### Impact of Farm

Not all subgroups in each clade showed the same pattern of geographic distribution, although some subgroups were exclusively comprised of isolates from a single farm. For example, all Subgroup A2 isolates were from Farm III and all Subgroup B1 isolates from Farm X. In contrast, Subgroups B3 only contained two from Farm I; A3 contained isolates from Farms V and VII.

Our phylogeny revealed that some isolates from different farms can be grouped together and were closely related: isolates in Subgroup A3 obtained from Farms V and VII with few to no SNP differences among them. For example, CFSAN051854 (CFSAN051854_spleen(m7)_1996_FarmV_SCA) and CFSAN051864 (CFSAN051864_spleen_1996_FarmVII_SCA) where zero SNP differences were observed (Table S1).

Isolates from some farms were only distantly related and, unsurprisingly, our phylogeny showed these belonging to different subgroups. For example, Subgroups B1 and B2 both contained isolates from Farm X, indicating that these were distantly related to the rest of the isolates from our sequencing.

There were several cases in which isolates from different farms were found to belong to the same subgroup: isolates from Subgroup B2, which contained 11 clade-defining SNPs, came from Farms VI, X, and XI. Isolates in Subgroup A3 were found at Farms V and VII, and there were only very small differences among their SNPs (Table S1).

Although isolates in Subgroups A1 and A5 were found at different farms, isolates from the same farm shared common ancestors. Specifically, all Subgroup A1 isolates were from Farm XII and XV, isolates from Farm XII formed a cluster and shared a common ancestor, and another common ancestor was shared by all isolates collected from Farm XV.

#### Impact of Isolation Year

Isolates in each subgroup were collected during the same year, with only two exceptions: A1 contained isolates from 1995 and 1996, and A5 contained isolates from 1997 and 1998. In Subgroup A1, isolates from 1995 were grouped together sharing common ancestor, which also applied to those from 1996 in Subgroup A1. In another case, all Subgroup A5 isolates were collected from 1998 except CFSAN063788, which was from 1997.

#### Impact of Isolation Organ

As expected, isolates from the same mouse appeared very closely related: SNP differences ranged from zero (m8) to two SNPs (m2). Most subgroups contained isolates from both organs. Although Subgroups A4, A8, B3, and B4 only contained isolates originating from spleens, our phylogenetic analyses did not reveal any organ-defining SNPs that could be reliably used to distinguish between SE isolates taken from spleens and those obtained from intestines.

### Pathogen Detection SNP Cluster analysis

At the time of this research (Dec 7^th^, 2017), the NCBI Pathogen Detection Isolates Browser (https://www.ncbi.nlm.nih.gov/pathogens) contained more than 94,000 *Salmonella enterica* genomes. At the time of our analysis, 86 of our isolates fit into five existing Pathogen Detection SNP Clusters, as follows. All 68 isolates, but CFSAN063803, within our eight Clade A subgroups belonged to one single Pathogen Detection SNP Cluster, which was designated as SNP Cluster A (SCA, at the time designated as SCA PDS000002757.323) (28). CFSAN063803 did not fit within any of the established SNP Cluster at that time. The four subgroups we recognized as Clade B belonged to four different Pathogen Detection SNP Clusters, which were designated as SCB, SCC, SCD, and SCE, respectively.

The data from Pathogen Detection Isolates Browser matched our phylogenetic analysis. Among our sequenced isolates, some farms contained isolates that were distantly related according to Pathogen Detection Isolates Browser data. For example, isolates collected from mice at Farm I, which we identified as Subgroups A8, B3, and B4, were members of three existing Pathogen Detection SNP Clusters: SCA, SCB, and SCC, respectively.

Our isolates in SCA had been collected from mice at 12 different farms between 1995 and 1998. However, SCA also encompassed 5,468 genomes already in the Pathogen Detection Isolates Browser. This provides the opportunity to explore additional levels of relatedness across SE isolates, as well as identify patterns across multiple years. For example, in the Pathogen Detection phylogenetic tree, Subgroup A1 isolates from 1995 shared a common ancestor with SE NYVetLIRN-37 (Sequence Read Archive (SRA) number: SRR6107632), which was isolated from dust taken from a poultry coop at Massachusetts in April 2017 (https://www.ncbi.nlm.nih.gov/Structure/tree/#!/tree/Salmonella/PDG000000002.1124/PDS000002757.351). Another example, SE WAPHL_SAL-A00192, which was isolated from an avian source from Washington in 2003, shared a common ancestor with Subgroup A5 isolates (https://www.ncbi.nlm.nih.gov/Structure/tree/#!/tree/Salmonella/PDG000000002.1124/PDS000002757.351).

It was notable that isolates from egg yolk and chicken drag swab appeared closely related to isolates in Subgroups A4 and A6. For example, SE CRJJGF_00137 (egg yolk, 2002, US, SRR1686612) and SE OH-10-18938-5 (chicken drag swab, 2010, Ohio, SRR5278942) were closely related to CFSAN051834 and CFSAN051835 in Subgroup A4 (https://www.ncbi.nlm.nih.gov/Structure/tree/#!/tree/Salmonella/PDG000000002.1124/PDS000002757.351).

The SCB (designated at that time as PDS000004690.16) encompassed a total of 24 isolates including those five isolates of our Subgroup B4. These 24 isolates in SCB were obtained from human, animal, food, and environmental sources in US and Canada (Figure 2). Within SCB, our Subgroup B4 isolates were clustered together and shared a most recent common ancestor with five NCBI isolates collected from human stool (SE PNUSAS011122, US, 2016), turkey (SE SA19943269, Canada), and chicken drag swab (SE OH-15-14655, OH, US, 2015, SE OH-12-29345, OH, US, 2012 & SE OH-13-28244, OH, US, 2013). The remaining 14 isolates in SCB formed a separate cluster, these were 13 clinical isolates and one environmental isolate that all shared a different common ancestor from the rest of SCB. The minimum distance between isolates in SCB was one SNP while the maximum number was 104.

**Figure 2.**
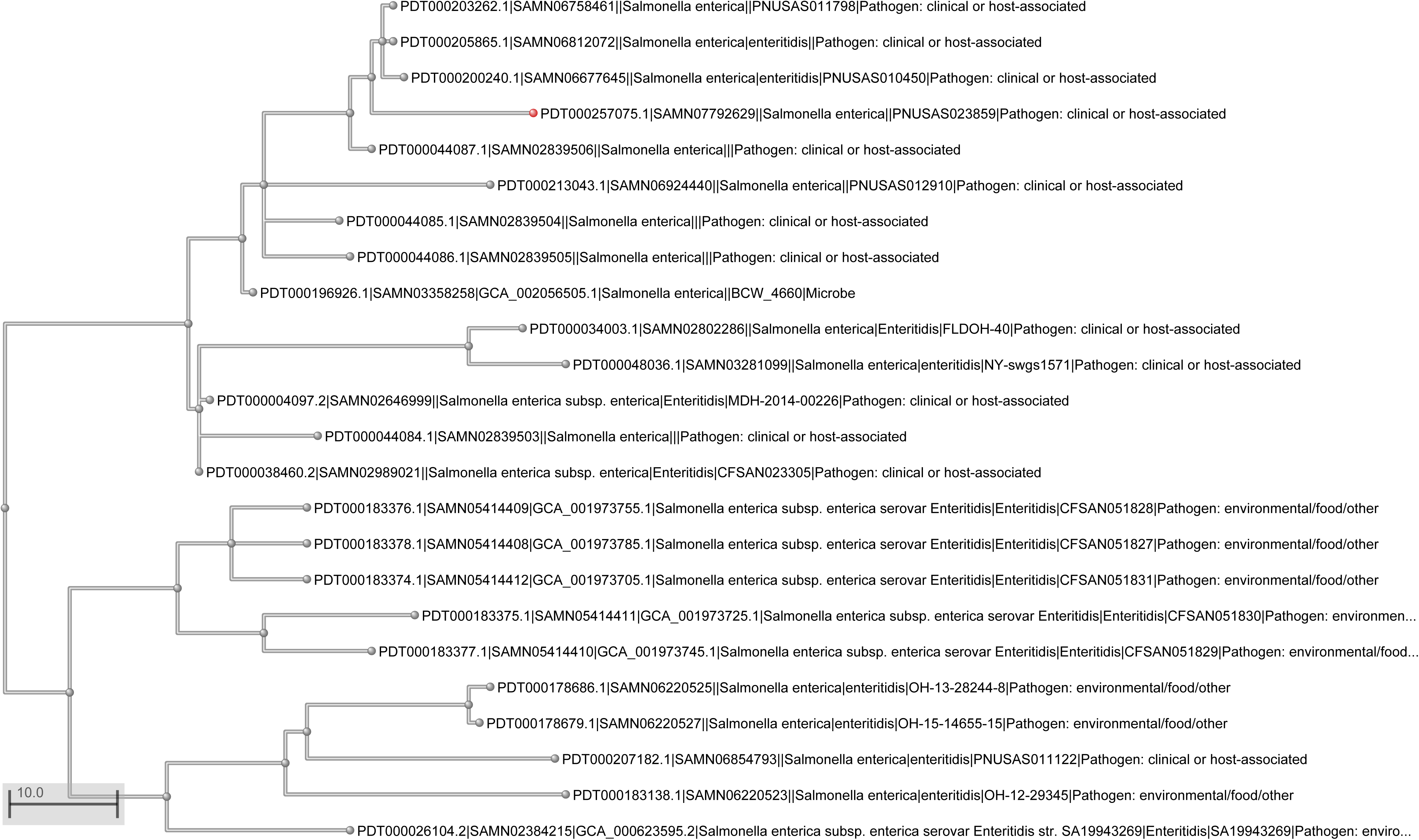
Phylogenetic tree of SNP Cluster B (SCB, designated at that time as PDS000004690.16) from NCBI Pathogen Detection Isolates Browser. The phylogenetic tree encompassed 24 isolates including our five sequenced isolates belonging to Subgroup B4.

SCC (designated at that time as PDS000011158.1) consisted of two isolates from Subgroup B3. No other genomes from Pathogen Detection Isolates Browser fit within SCC. Similarly, no other NCBI genome fit within SCD (designated at that time as PDS000011157.1), which contained only Subgroup B1 isolates. SCE (designated at that time as PDS000004693.11) comprised 20 isolates from chicken, mouse, and human. These isolates had been collected from the states of Tennessee, Georgia, and Pennsylvania, in the US. Eight of the Subgroup B2 isolates that fit within SCE shared a common ancestor (Figure 3). Intriguingly, all SCE isolates, with the exception isolate PNUSAS014592, carried at least one of following antimicrobial resistance genes: *tetA*, *aadA*, *bla*_*TEM*-*1*_ (https://www.ncbi.nlm.nih.gov/pathogens/isolates#/tree/Salmonella/PDG000000002.1056/PDS000004693.11/).

**Figure 3.**
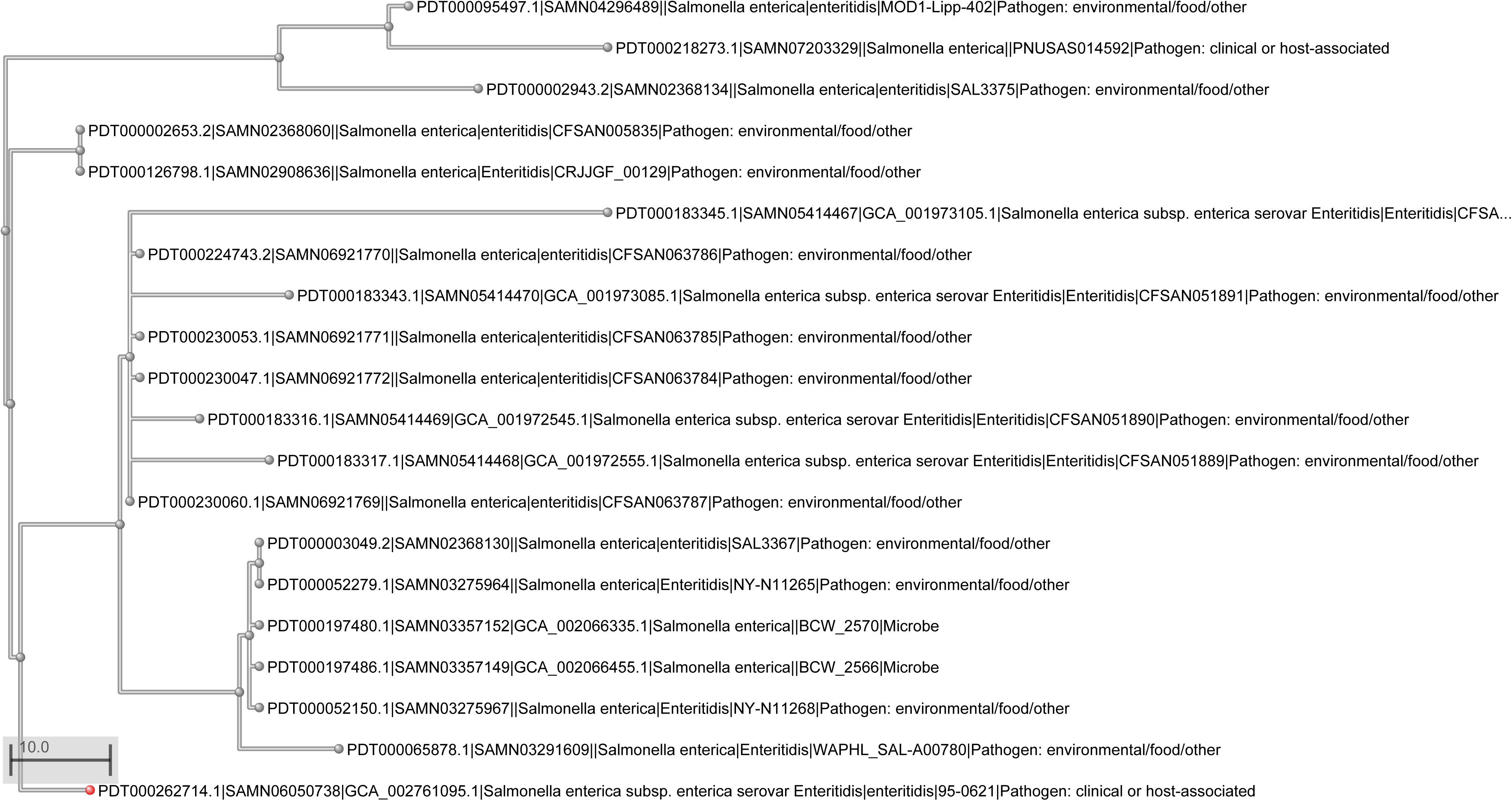
Phylogenetic tree of SNP Cluster E (SCE, designated at that time as PDS000004693.11) from NCBI Pathogen Detection Isolates Browser. The phylogenetic tree encompassed 20 isolates including our eight sequenced isolates belonging to Subgroup B2.

### Clade-defining SNPs

We identified clade-defining SNPs and annotations identifying synonymous/nonsynonymous changes in amino acids, positions in reference genes, strands, and gene functions are presented in Table 3.

**Table 3.**
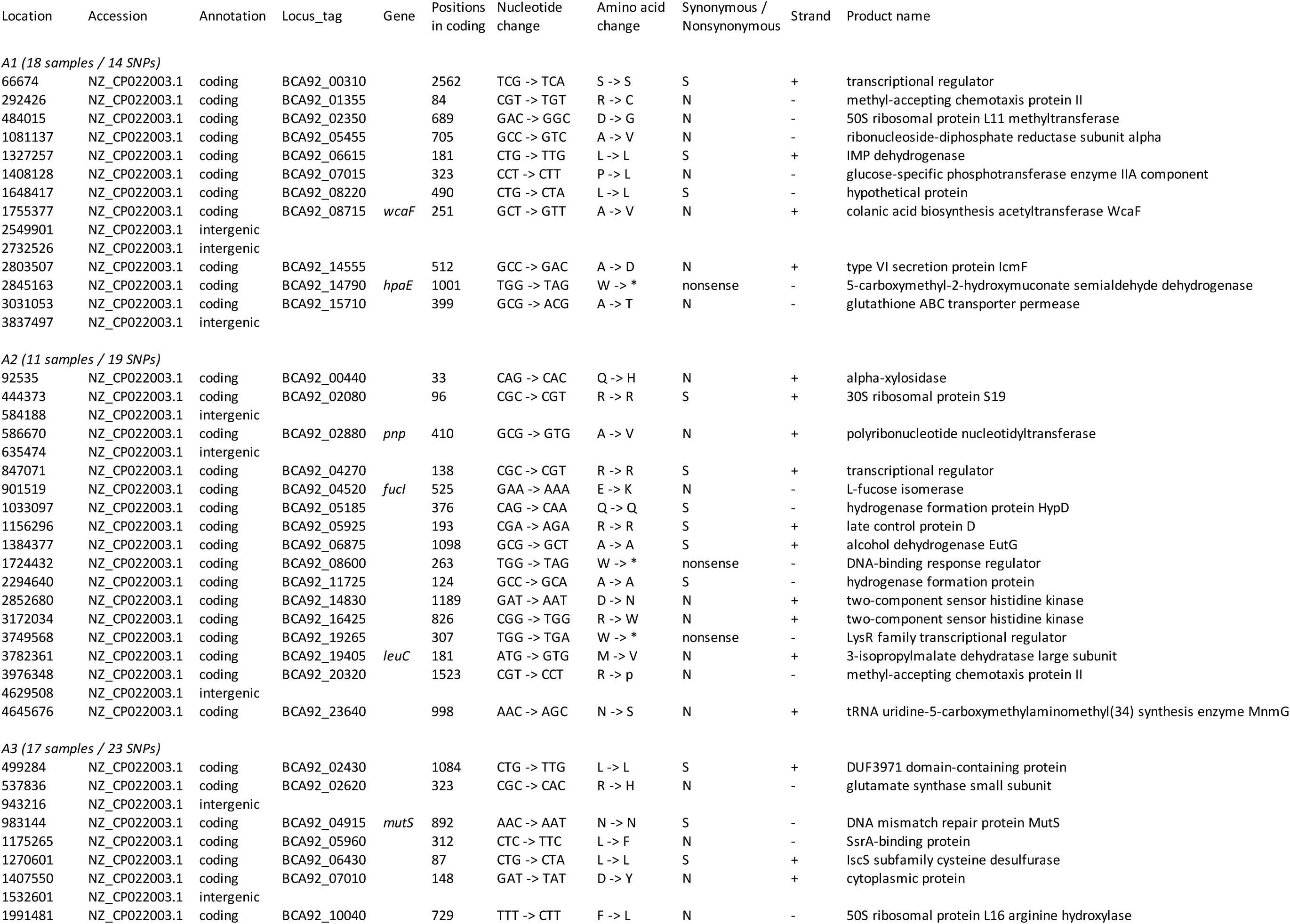
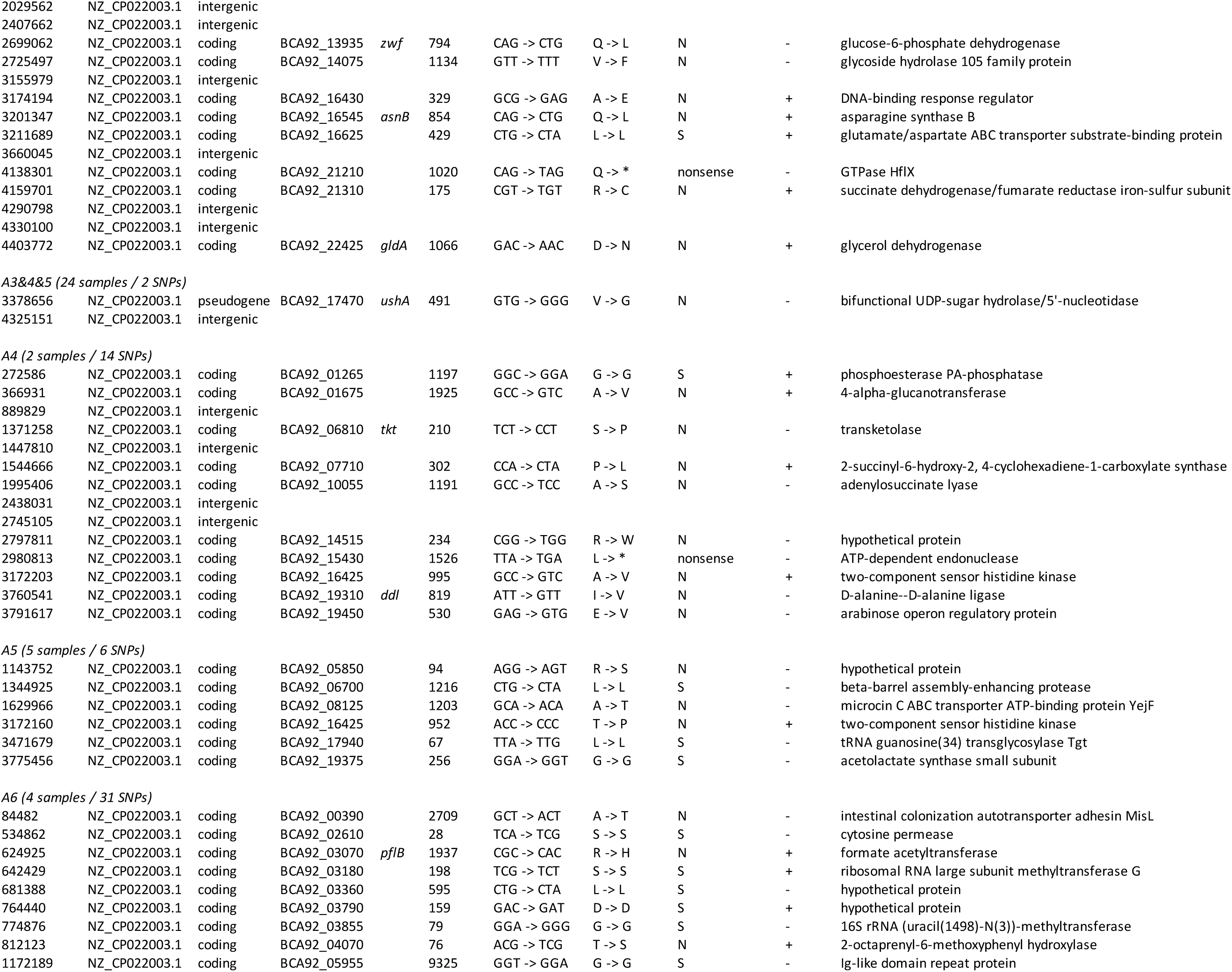
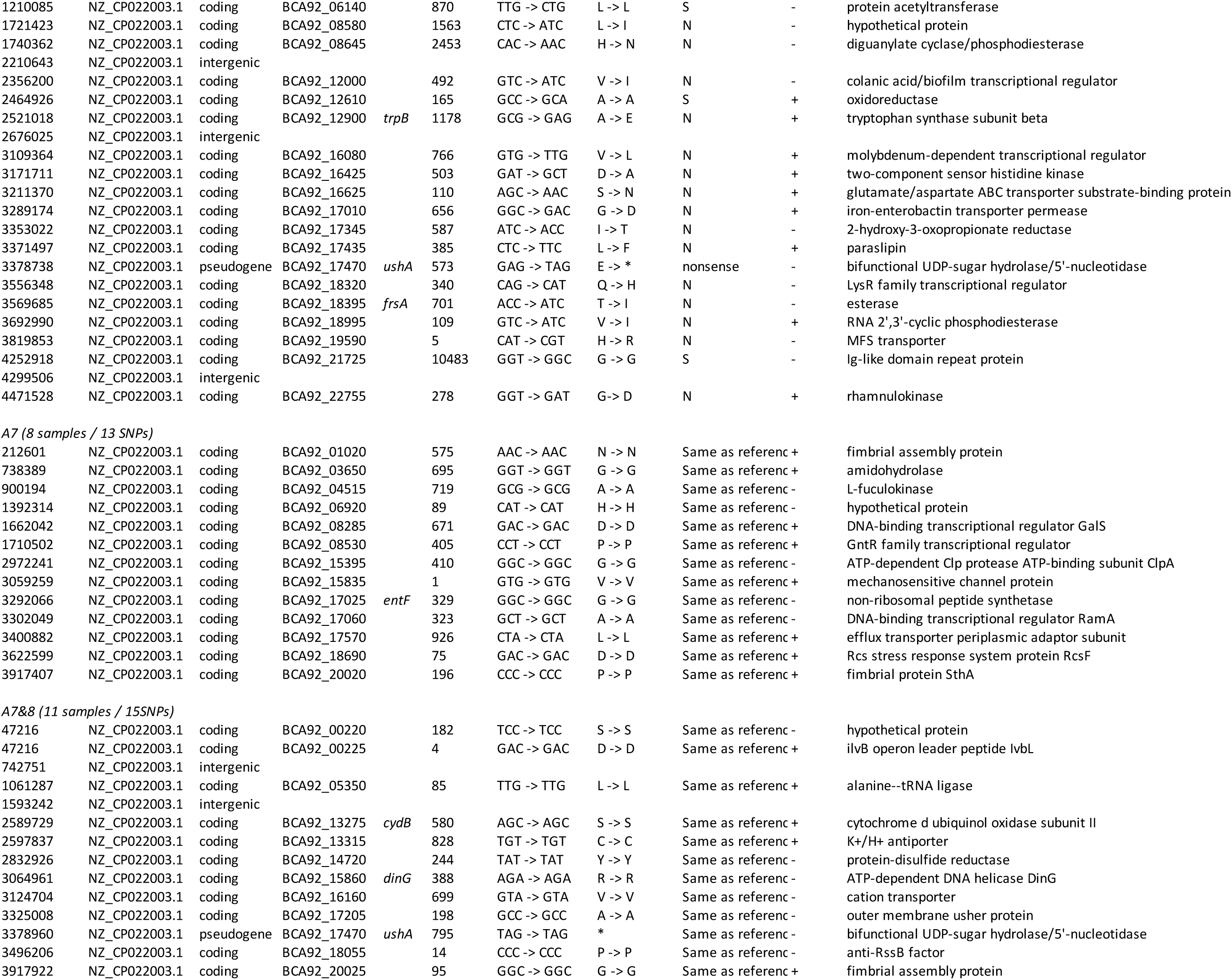
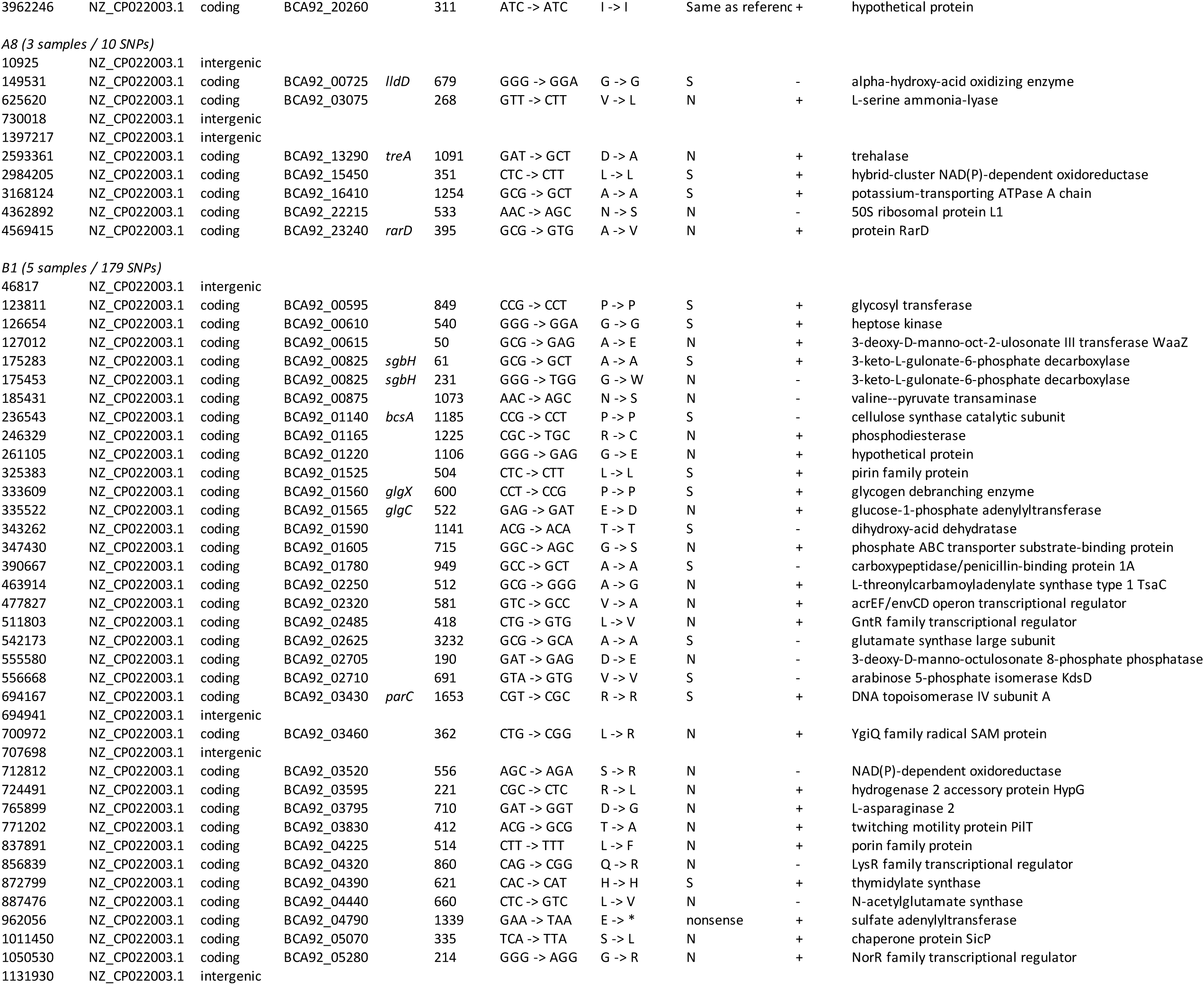
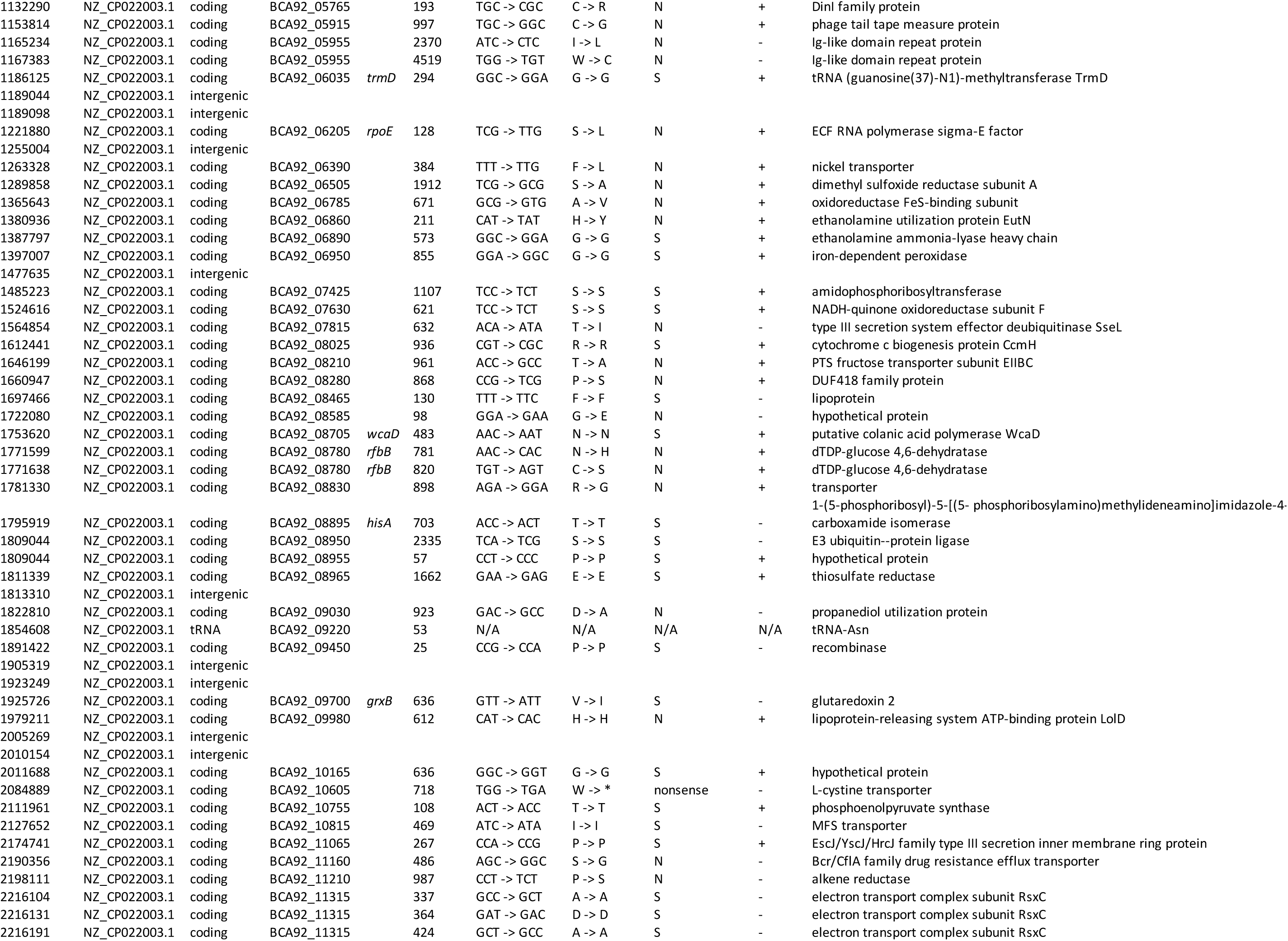
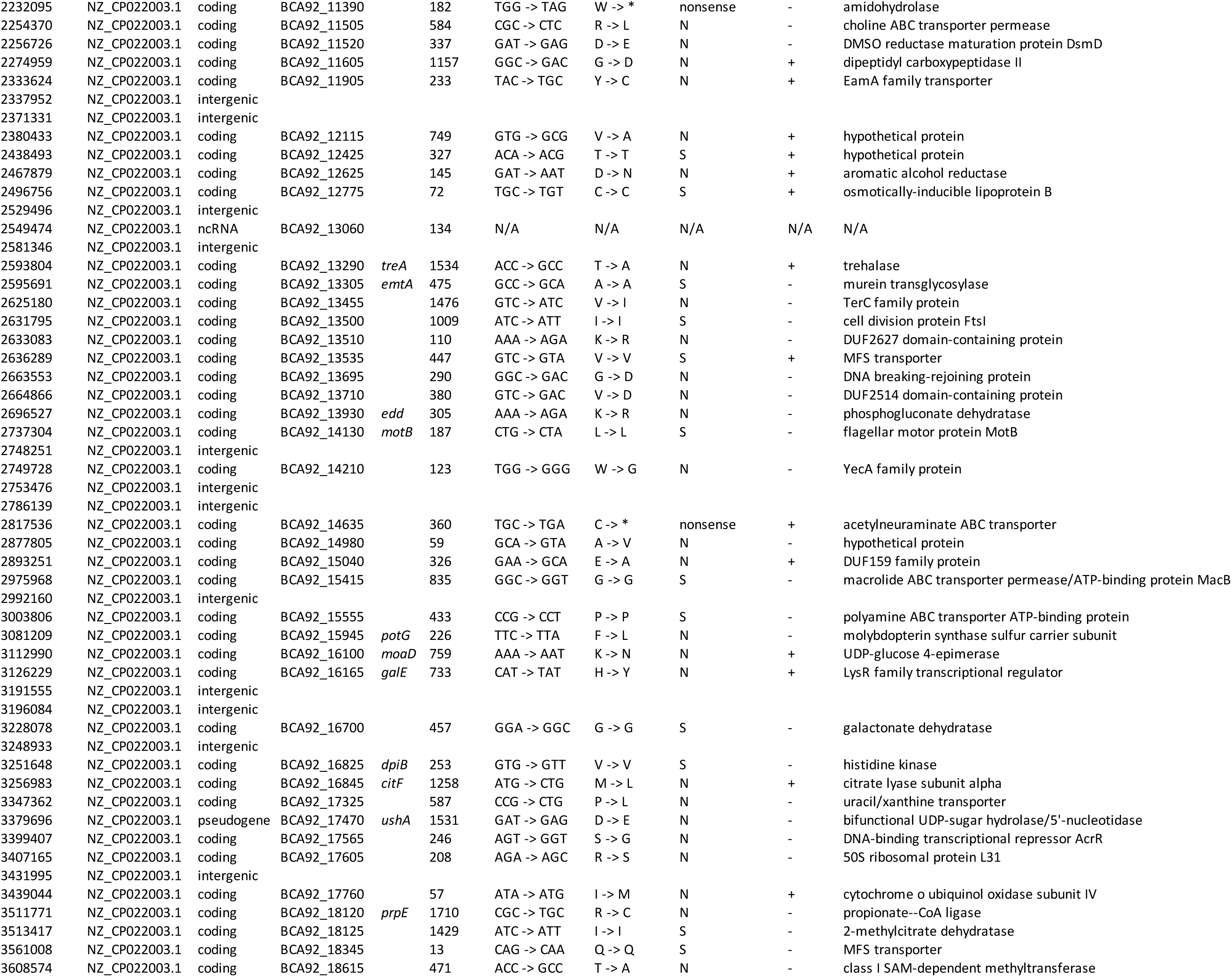
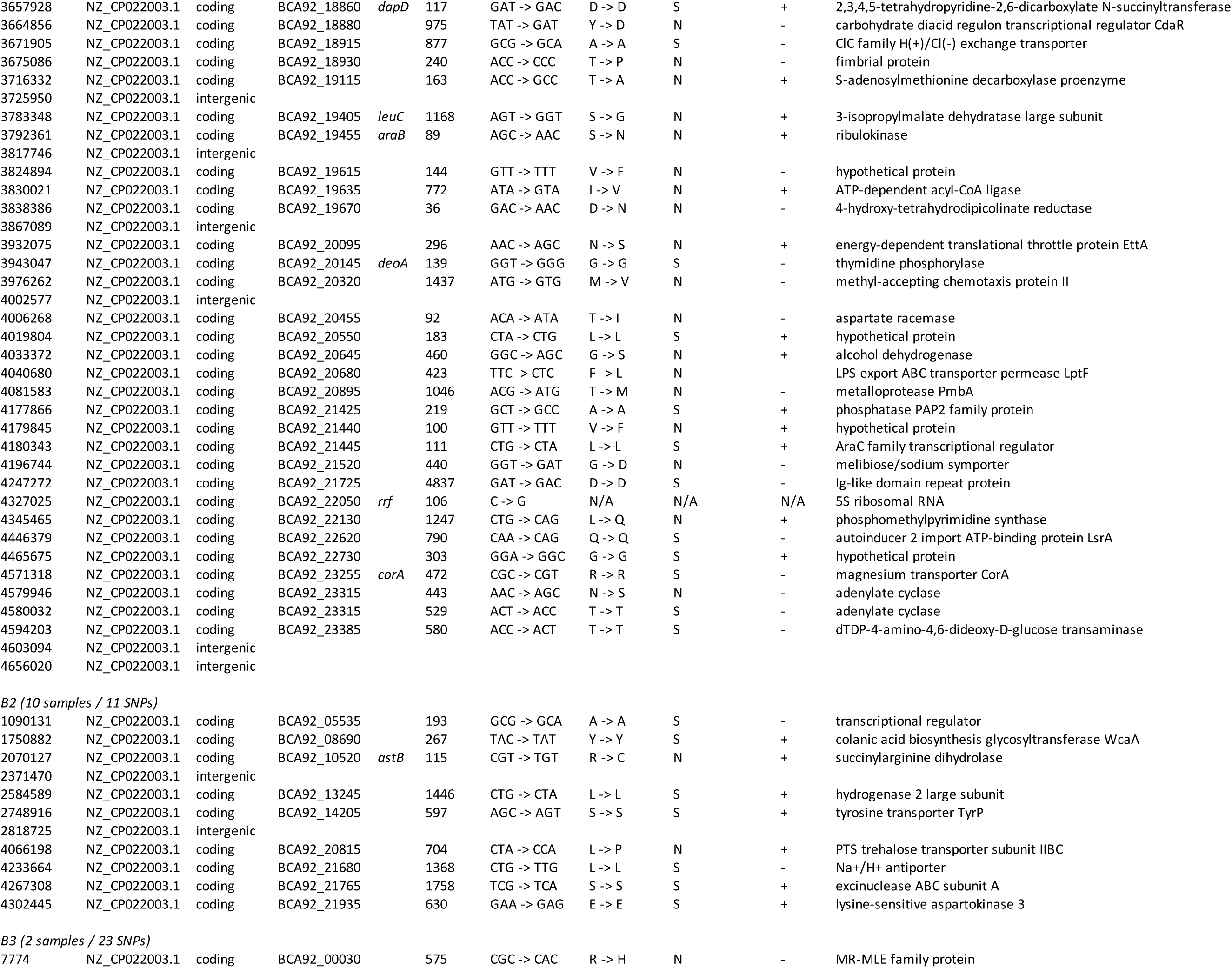
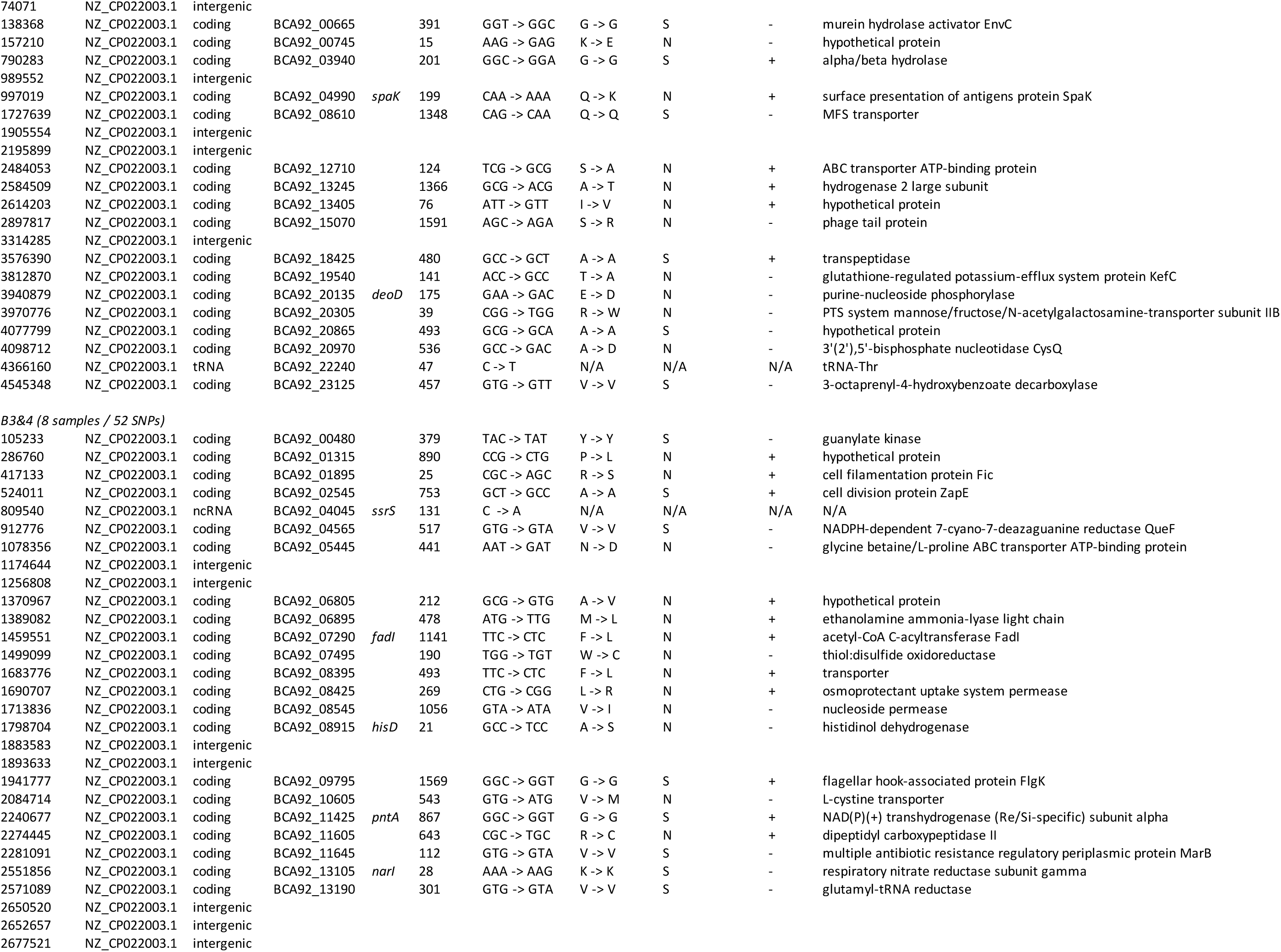
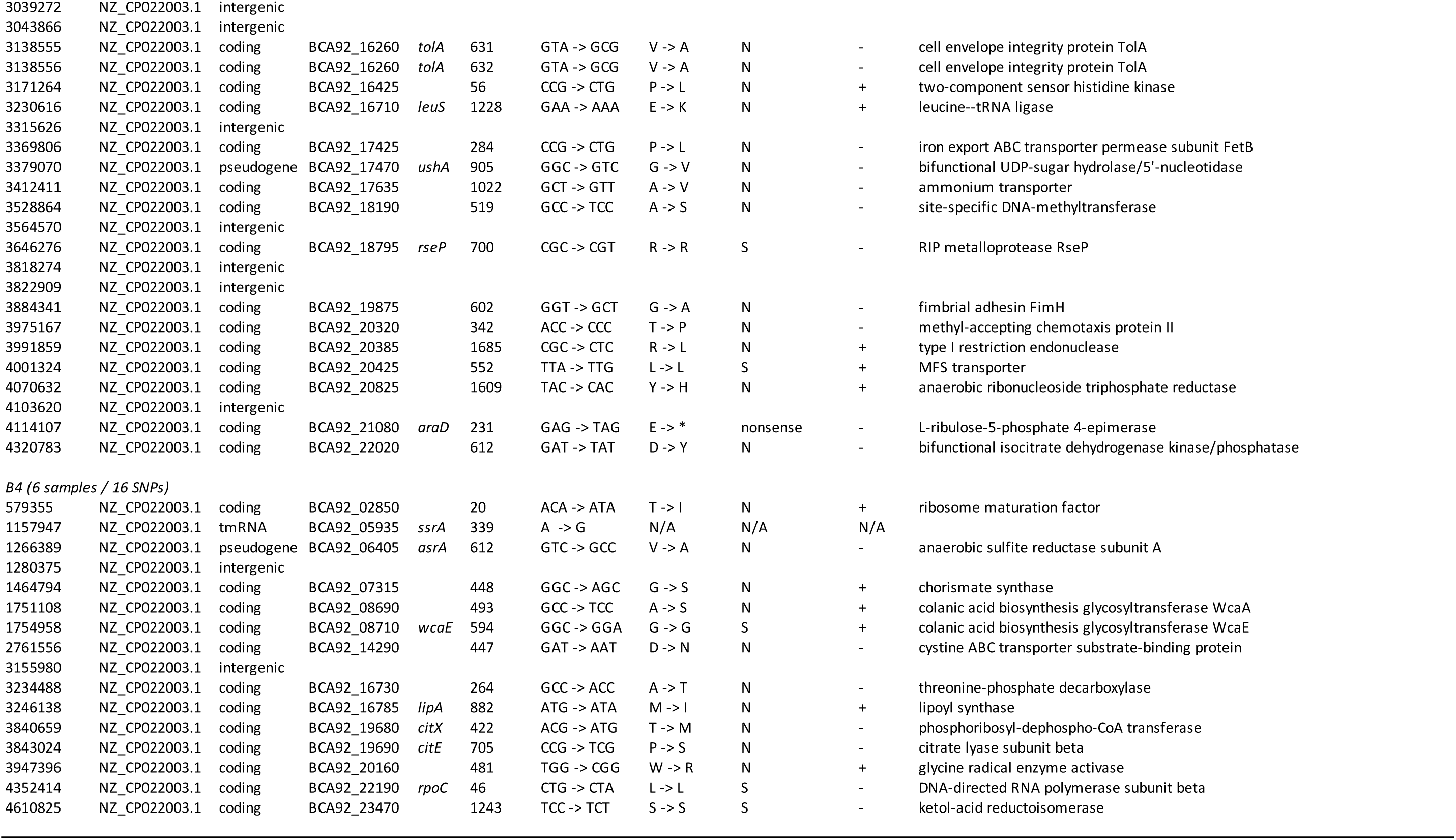
Variable genes observed that define subgroups in phylogenetic tree of S. Enteritidis

#### Clade A polymorphisms

We identified 11 SNPs that defined Subgroup A1, including seven nonsynonymous changes, three synonymous changes, and one nonsense mutation. Type VI secretion protein IcmF (reference locus tag BCA92_14555) contained one C to A mutation, which resulted in amino acid changing A to D. Another unique genetic signature change within Subgroup A1 occurred in the colanic acid synthesis gene *wcaF* (BCA92_08715), which changed C to T change. The nonsense mutation resulted in a stop codon which interrupted *hpaE* (BCA92_14790), encoding for enzymes involved in catabolism in the aromatic pathway.

In Subgroup A2, which contained isolates exclusively from Farm III, we discovered 19 clade-defining SNPs, including 16 in coding region. The LysR family transcriptional regulator (dBCA92_19265) contained one G to A mutation resulting in a stop codon.

Other notable findings in other subgroups included nonsynonymous mutations in *zwf* (Subgroup A3, BCA92_10040, oxidoreductase in glucose metabolism), *asnB* (Subgroup A3, BCA92_16545, asparagine synthase B), *ushA* (Subgroups A4&A5 and A6, BCA92_17470, 5’-nucleotidase), and *frsA* (Subgroup A6, BCA92_18395, esterase).

#### Clade B polymorphisms

Among the 179 clade-defining SNPs in Subgroup B1, 146 SNPs were in coding regions, including 85 nonsynonymous mutations and four nonsense mutations. Subgroup B2, which contained isolates carrying resistance genes, contained 11 SNPs with nine in coding regions. Among the isolates in Subgroups B3 and B4, we identified multiple nonsynonymous mutations, including *deoD* (BCA92_20135, purine-nucleoside phosphorylase), *cysQ* (BCA92_20970, 3’(2’),5’-bisphosphate nucleotidase activity and magnesium ion binding), *hisD* (BCA92_08915, histidinol dehydrogenase and zinc ion binding), *tolA* (BCA92_16260, cell envelope integrity protein in transporter activity), and *fimH* (BCA92_19875, fimbrial adhesion).

## Discussion

The dissemination of SE via mice, particularly on poultry farms, is considered to be one of the most serious threats to poultry industry today (2). Here, we characterized a set of 91 SE that (i) represented two organs in mice that have been associated with dissemination of SE among poultry and hence to humans, (ii) were isolated at 15 farms in Pennsylvania during the mid-1990s, which was a time during which few SE isolates from mice have previously been sequenced, and (iii) analyzed in combination with the open access NCBI Pathogen Detection Isolates Browser. These steps allow us to construct a more nuanced picture of SE dissemination during the 1990s, and also identify connections between historic isolates and current SE phenotypes. Our study demonstrated that WGS not only reliably distinguishes among closely related SE isolates from mice and trace a genome back to its farm of origin and year of isolation, but also allows sufficient resolution to distinguish between SE isolates, even those collected from different organs (spleens and intestines) of individual mouse. In addition, our analyses showed that (i) isolates carrying antimicrobial resistance genes formed a separate subgroup, which could indicate a shared mechanism which enables that feature, (ii) open access WGS database contributes comprehensive perspectives to our understanding of selected isolates, and (iii) new clade-defining markers and NCBI Pathogen Detection Isolates Browser SNP Clusters were identified, offering tool with high resolution in outbreak investigations and rapid detections to identify specific clade related to certain years or locations.

Our results strongly suggested it was possible for unique ecologies of SE to develop on individual farms, although local adaptation is not inevitable. Farms I and X exemplify this range of possibilities: Farm I exhibited heterogeneous isolates while isolates from Farm X were shown to be highly similar. Isolates can spread from one location to another in multiple ways: insects (29), wild birds (30), wild animals (31, 32), and even wind (33) can move contamination from one place to another. However, among these possible transmission routes, mice are ubiquitous pests (8–10), and their behaviors may help shape those unique local ecologies: mice migrate periodically and also defend their territories. Understanding the genetic relatedness among the SE carried by mice and the SE found in veterinary, food, and human sampling will help improve safety and security in poultry industry.

### WGS data identified a subgroup consisting exclusively of isolates carrying antimicrobial resistance genes

Previously, WGS has been used to differentiate drug-resistant *S. enterica* isolates from different locations, which can exhibit notable differences in resistant-relevant genotypic and phenotypic characteristics (34). Other research has shown WGS can be valuable in predicting phenotypic resistance among both *S. enterica* (34, 35) and *E. coli* (36). In the current study, WGS analyses revealed that all our Subgroup B2 isolates carried *bla*_*TEM*-*1*_ and *tetA.* It is possible that Subgroup B2 isolates share specific genetic features that permit them to obtain and carry antimicrobial resistance genes via horizontal gene transfer, or make it more likely for those genes to be maintained. For example, bacteria that carry non-functional Clustered Regularly Interspaced Short Palindromic Repeats (CRISPR) /cas system could acquire plasmids carrying antimicrobial resistance genes. Possession of a fully-functioning CRISPR/cas system is reversely correlated with antimicrobial resistance in bacteria (37–39).

### Open access genome databases allow greatly expanded genomic and phylogenetic investigations

In the NCBI Pathogen Detection Isolates Browser, comprehensive data was available for each genome, including up to 40 columns of detail such as WGS run qualities, outbreak relatedness, and antimicrobial resistance genotypes. The Browser also assigns specific cluster ID numbers computed based on SNP distances. Although these cluster numbers can change as new information is added to the Browser, this feature allows researchers to quickly identify isolates most closely related to target isolates, which can assist in recognizing possible connections among clinical illness cases. The phylogenetic analyses from the Browser were consistent with our phylogenetic tree. Multiple subgroups in the current study formed distinct SNP Clusters containing isolates exclusively from our collection, like B3 isolates in SCC. We identified clinical isolates and poultry related isolates closely related to our isolates, such as SE PNUSAS011122 (human stool, US, 2016) and SE OH-15-14655 (chicken drag swab, OH, US, 2015) with B4 isolates in SCB. Our data has the potential to bridge surveillance data with long-term and large-scale genomics and phylogenetics studies (19).

### Genetic variations in clade-defining SNPs showed possible unique genotypic and phenotypic features

Distinctive genetic features are extremely useful for epidemiologic investigations. Finding such genetic identifiers can help rapidly determine outbreak lineages and accurately distinguish highly clonal clades (6). The nonsynonymous changes we identified in this study suggested that a combination of several genetic factors has facilitated the survival and growth of SE, resulting in different contamination risks for each subgroup. For example, the *icmF* we identified in Subgroup A1 was part of Type VI Secretion System, which is known to be required for full virulence in mice (40, 41). Similarly, *fimH* alleles have been associated with the abilities of *Salmonella* to bind onto avian or mammalian cells (42). Despite the clonal structure of SE, isolates vary greatly in the ability to contaminate eggs, which is biologically independent of phage types those isolates belong to (15, 43, 44). The heterogeneity of metabolic profiles in SE isolates might provide an explanation for the variation in contamination capability (15). The accumulation of mutations that affect gene function is a significant part of the process by which *S. enterica* becomes host adapted (45). Such host adaptations may well be occurring at some of the farms where we collected SE from local mice, with important consequences for the safety and security of the poultry supply chain. Notably, serovars Enteritidis, Gallinarum, and Pullorum can circulate within the same farm, and sometimes within the same bird, as evidenced by field analyses conducted in South America (46). Therefore, WGS also has potential for detecting evolutionary trends within SE that could threaten the poultry industry supply chain. Our data also pave the way for research on poultry pathogenic serovars S. Gallinarum and S. Pullorum, which diverged independently from an Enteritidis-like ancestor (3, 47, 48).

## Acknowledgements

The authors thank Dr. Lili Fox Vélez for scientific writing support and Oak Ridge Institute for Science and Education (ORISE) Research Participation Program at the U.S. Food and Drug Administration support on this project.

